# Mutation accumulation in chromosomal inversions maintains wing pattern polymorphism in a butterfly

**DOI:** 10.1101/736504

**Authors:** Paul Jay, Mathieu Chouteau, Annabel Whibley, Héloïse Bastide, Violaine Llaurens, Hugues Parrinello, Mathieu Joron

## Abstract

While natural selection favours the fittest genotype, polymorphisms are maintained over evolutionary timescales in numerous species. Why these long-lived polymorphisms are often associated with chromosomal rearrangements remains obscure. Combining genome assemblies, population genomic analyses, and fitness assays, we studied the factors maintaining multiple mimetic morphs in the butterfly *Heliconius numata*. We show that the polymorphism is maintained because three chromosomal inversions controlling wing patterns express a recessive mutational load, which prevents their fixation despite their ecological advantage. Since inversions suppress recombination and hamper genetic purging, their formation fostered the capture and accumulation of deleterious variants. This suggests that many complex polymorphisms, instead of representing adaptations to the existence of alternative ecological optima, could be maintained primarily because chromosomal rearrangements are prone to carrying recessive harmful mutations.

---

Polymorphic complex traits, which implicate the coordination of multiple elements of phenotype, are often controlled by special genetic architectures involving chromosomal rearrangements. Examples include dimorphic social organization in several ant species (1), coloration and behavioral polymorphisms in many birds and butterflies (2–6), dimorphic flower morphology in plants (7), as well as the extreme cases provided by sexual dimorphism encoded by the extensively re-arranged sex chromosomes. Why these polymorphisms arise is a long-standing question in biology (8–12).

The so-called supergenes controlling these striking polymorphisms are characterized by the suppression of recombination between linked loci, often through polymorphic chromosomal rearrangements which are thought to preserve alternative combinations of co-adapted alleles (1, 4, 5, 7, 12). The encoded phenotypes are often assumed to reflect the existence of multiple, distinct adaptive optima, and are frequently associated with antagonistic ecological factors such as differential survival or mating success (3, 13–15). Yet why and how alternative chromosomal forms become associated with complex life-history variation and ecological trade-offs is not understood.

The Amazonian butterfly *Heliconius numata* displays wing pattern polymorphism with up to seven morphs coexisting within a single locality, each one engaged in warning color mimicry with distinct groups of toxic species. Adult morphs vary in mimicry protection against predators and in mating success via disassortative mate preferences (13, 16). Polymorphic inversions at the mimicry locus on chromosome 15 (supergene P) form three distinct haplotypes (5). The standard, ancestral haplotype constitutes the class of recessive P alleles and is found, for example, in the widespread morph *silvana*. Two classes of derived haplotypes are known, both associated with a chromosomal inversion called P_1_ (∼400kb, 21 genes), each conferring increased protection against predatory attacks via mimicry. The first derived haplotype, encoding the morph *bicoloratus*, carries P_1_ alone; the second class of derived haplotypes carries P_1_ linked with additional yet still uncharacterized rearrangements (called BP2 in (5)) and occurs in morphs which typically exhibit intermediate levels of dominance, such as *tarapotensis* and *arcuella*. Inversion polymorphism and supergene formation originated via the introgression of P_1_ from the *H. pardalinus* lineage (17). The series of chromosomal rearrangements initiated by introgression allows us to unravel the stepwise process by which structural variation has become associated with directional and balancing selection.

Comparative analysis of *de novo* genome assemblies of 12 *H. numata* individuals revealed a history of supergene formation characterized by the sequential accretion of three adjacent inversions with breakpoint reuse. Pairwise alignment of assemblies shows that all derived haplotypes belonging to the intermediate dominant allelic class display two newly-described inversions: P_2_ (200kb, 15 genes), adjacent to P_1_, and the longer P_3_ (1150 kb, 71 genes), adjacent to P_2_ (Fig 1A, Sup. Fig. S1, Sup. Fig. S2). Sliding-window PCA along the supergene region confirmed the dominance of derived arrangements (denoted Hn1 and Hn123) to the ancestral arrangement (denoted Hn0) and their prevalence across all populations of the Amazon (Fig 1B, Fig 1C, Sup. Fig. S3, Sup. Fig. S4). Multiple genes in the inverted regions showed significant differential expression compared to ancestral segments, but this likely reflects divergence rather than direct breakpoint effects (Sup. Fig. S5). Indeed, none of the break-points of P_1_, P_2_ or P_3_ fell within a gene, and no transcript found in Hn0 specimens was missing, disrupted, or differentially spliced in specimens with inversions (Hn1 and Hn123).

**Fig. 1.**
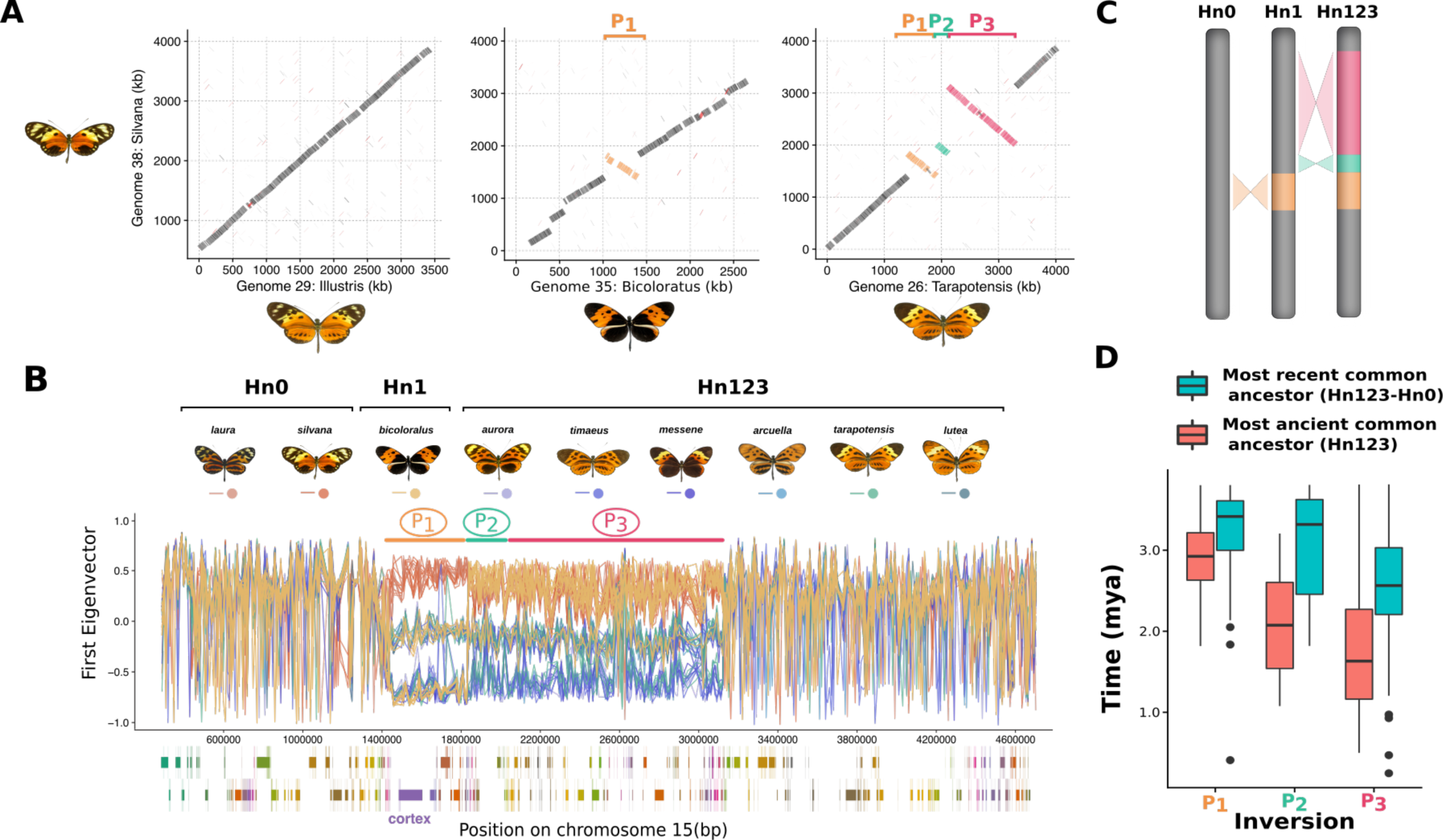
Genomic architecture of the *H. numata* wing pattern polymorphism. **A.** Alignment of the genome assemblies from 4 *H. numata* morphs across the supergene region on chromosome 15. **B.** Sliding window Principal Component Analysis (PCA) computed along the supergene (non-overlapping 5kb windows). For clarity, only a subset of morphs are shown here (full dataset presented in Sup. Fig. S3). Each colored line represents the variation in the position of a specimen on the first PCA axis along chromosome 15. Within the inversions, individual genomes are characterized by one of three genotypes : homozygous for the inversion (down), heterozygous (middle), homozygous for the standard arrangement (top). The gene annotation track is shown under the plot, with the forward strand in the lower panel and the reverse strand in the upper panel. Each gene is represented by a different colour **C.** Structure of the *H. numata* supergene P. Three chromosome types are found in *H. numata* populations, carrying the ancestral gene order (Hn0), inversion P_1_ (Hn1), or inversions P_1_, P_2_ and P_3_ (Hn123). **D.** Analysis of divergence times between Hn123 and Hn0 at inversions segment. The TMRCA between Hn123 and Hn0 and the most ancient common ancestor of Hn123 provide respectively the upper and lower bound of the inversions formation time. Boxplots display the distribution of estimated times computed on 5kb sliding window across the supergene (estimates plotted along the supergene presented in Sup. Fig. S7).Time intervals are consistent with the stepwise accretion of P_1_, P_2_ and P_3_, but the simultaneous origin of P_2_ and P_3_ cannot be formally rejected.

In contrast to the introgressive origin of P_1_(Sup Fig. S6, (17)), inversions P_2_ and P_3_ are younger and originated within the *H. numata* lineage. Upper and lower estimates of inversion ages, obtained by determining the most recent coalescence events between Hn0+Hn1 and Hn123, and within Hn123, respectively, suggest that the P supergene has evolved in three steps, involving the introgression of P_1_followed by the successive occurrence of P_2_ and P_3_ between ca. 1.8 and 3.0 Mya (Fig. 1D, Sup. Fig. S7). Haplotypes show sizeable peaks of differentiation (Fst) across inversion blocks (Sup. Fig. S8), reflecting their distinct histories of recombination suppression and confirming the stepwise accretion of these inversions. The three adjacent inversions underlying the mimicry polymorphism of *H. numata* are therefore of distinct ages and originated in distinct lineages, which provides an opportunity to partition their mutational history and distinguish the consequences of their formation from those resulting from their maintenance in a polymorphism.

Since chromosomal regions carrying inversions rarely form chiasma during meiosis, recombination is strongly reduced among haplotypes with opposite orientations (18). Recombination suppression between structural alleles is predicted to lead to inefficient purging of deleterious variants and therefore to the accumulation of deleterious mutations and transposable elements (TEs) (19). Consistent with this prediction, estimation of the TE dynamics obtained by computing whole genome TE divergence supports a recent burst of TE insertion within the inversions, reported particularly by TEs belonging to the RC, DNA and LINE classes (Fig. 2A, Fig. 2B). Inverted haplotypes show a significant size increase (mean=+9.47%) compared to their corresponding non-inverted region in Hn0 (Fig. 2C) and this expansion was caused primarily (71.8%, Fig. 2A) by recent TE insertions from these classes (Fig. 2B, Sup. Fig. S9).

**Fig. 2.**
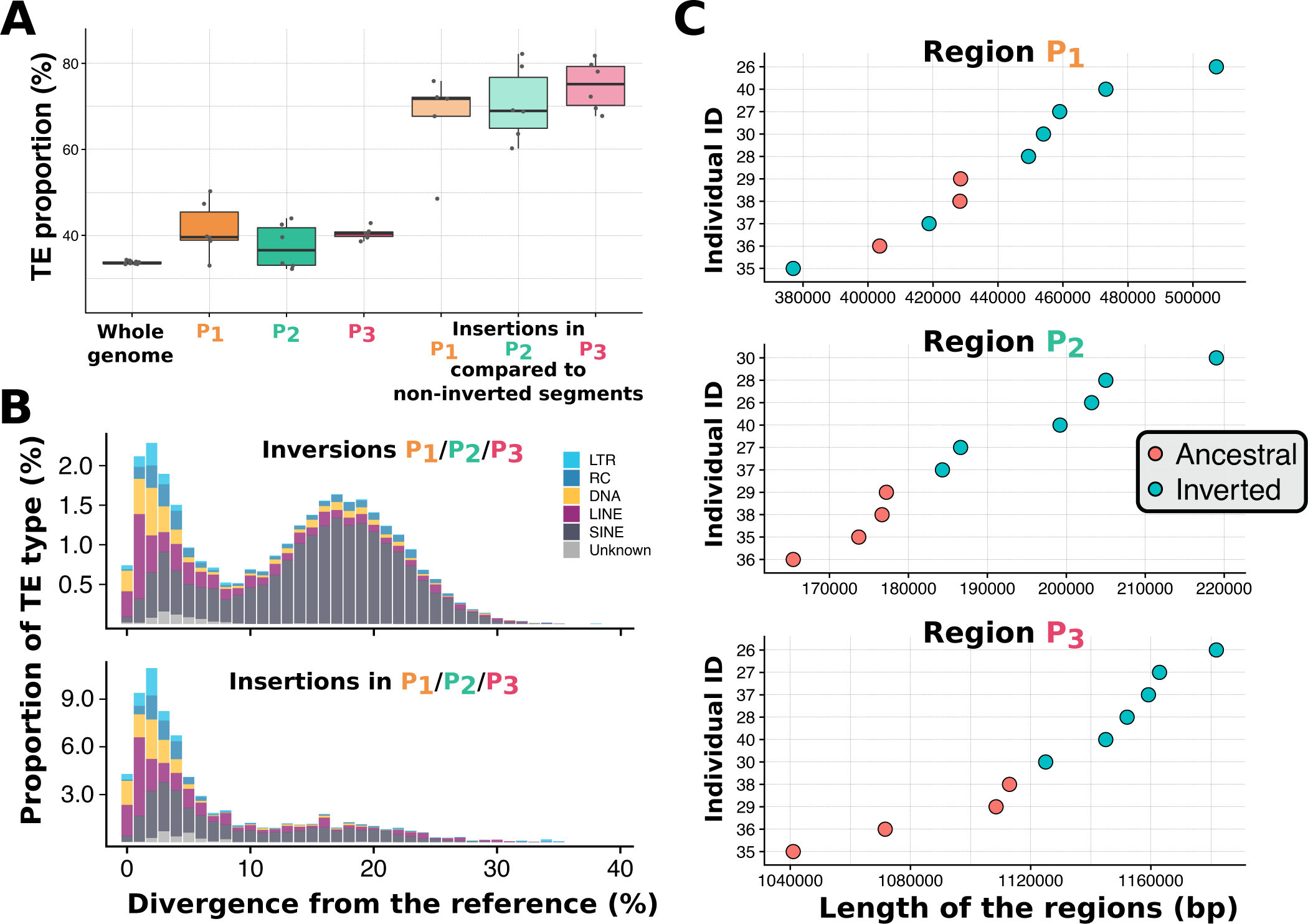
Variation in inversion size due to accumulation of transposable elements. **A.** Proportion of transposable elements in the whole genome, in the 3 inversions, and in the region present uniquely in inversion P_1_, P_2_ or P_3_ and not in ancestral non-inverted haplotype -i.e. sequences that were inserted in P_1_, P_2_, or P_3_. Insertions in inversions are mostly transposable elements. **B.** Timing of insertion, in units of nucleotide divergence, for the distinct classes of transposable elements found in inversions or only in sequences that were inserted in P_1_, P_2_, or P_3_. Recently active TEs (RC, DNA and LINE) are those that have accumulated within inversions. **C.** Size comparisons of orthologous standard and inverted chromosomal segments. Inverted haplotypes are longer than haplotypes with the ancestral gene order.

To investigate the impact of polymorphic inversions on the accumulation of deleterious mutations, we calculated, independently on inverted and non-inverted segments, the rate of non-synonymous to synonymous polymorphism (pN/pS), the rate of non-synonymous to synonymous substitution (dN/dS) and the direction of selection (DoS, (20)). Consistent with a low efficiency of selection in eliminating deleterious variants, P_1_, P_2_, and P_3_ were all found to be enriched in non-synonymous relative to synonymous polymorphisms compared to the whole genome and to non-inverted ancestral segments (pN/pS_P1_=0.83, pN/pS_P2_=0.54, pN/pS_P3_=0.49, Fig. 3A, Sup. Tab. S12). The inversions were also found to be under negative selection (DoS_P1_=-0.136, DoS_P2_ = −0.087, DoS_P3_=-0.079), with values reflecting their sequential origin (Fig. 3A, Sup. Tab. S12). Because P_1_ was introgressed from the *H. pardalinus* lineage (Sup. Fig. S6, (17)), mutations that accumulated in P_1_ before the introgression (i.e. shared with *H. pardalinus*) could be distinguished from those arising after supergene formation in *H. numata* (i.e. unique to Hn1 and Hn123). This revealed that non-synonymous mutations which existed in the P_1_ segment before the introgression underwent a high rate of fixation in *H. pardalinus* (dN/dS = 0.78, Sup. Fig. S10), and in *H. numata* (dN/dS=1.33, Fig 3B), suggesting that both the formation of P_1_ and its introgression led to the fixation of deleterious mutations. By contrast, 99.9 % of the mutations that accumulated in coding regions of P_1_ after its introgression -i.e. after super-gene formation-remain polymorphic in Hn1/Hn123 and a high proportion of them are non-synonymous (dN/dS=0.00, pN/pS=0.978, DoS=-0.49, Fig 3B, Sup. Tab. S12). Taken together, these results suggest that the inversions have captured and accumulated deleterious mutations during their evolution, presumably owing to bottlenecks generated by their formation and to recombination suppression with their ancestral, coexisting counterparts

**Fig. 3.**
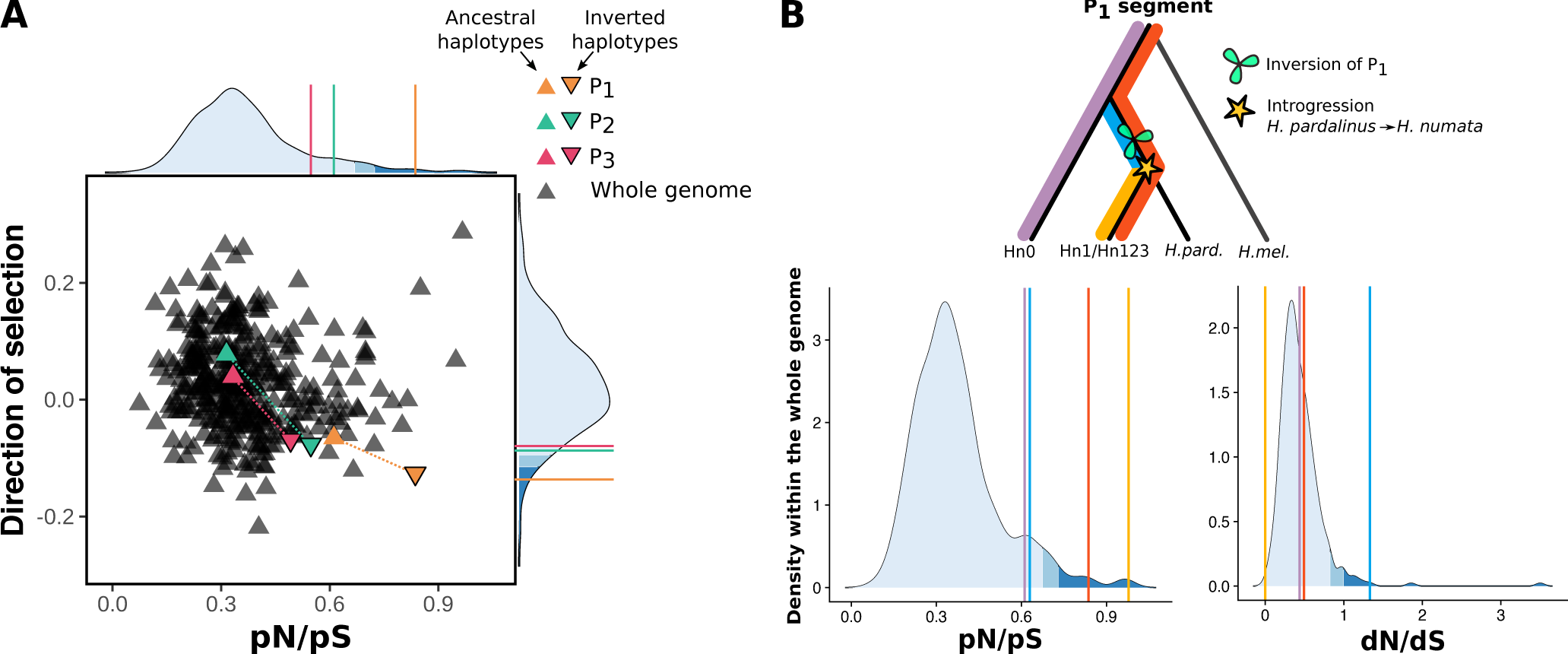
Accumulation of deleterious variants in inversions. **A.** Direction of selection and ratio of non-synonymous to synonymous polymorphisms (pN/pS) ratio, computed on 500 kb windows genome-wide and in the inversions segments, for both inverted and non-inverted haplotypes. Only genes with coding sequences >5kb (n=6364) were retained in this analysis. Inversions tend to be under negative selection and to accumulate non-synonymous polymorphism. **B.** Ratios of non-synonymous to synonymous substitutions (dN/dS) and polymorphisms (pN/pS) on the different mutations partitions observed in the P_1_segment : all mutations observed in Hn0 (purple), all mutations observed in Hn1/Hn123 (red), all mutations shared by *H. pardalinus* and Hn1/Hn123 and not observed in Hn0 (blue) and all mutations present uniquely in Hn1/Hn123 (yellow).

Inversions with an accumulated mutational load are expected to incur a fitness cost. Indeed, *H. numata* inversions were found to have detrimental effects on larval survival in homozygotes. When comparing survival among P genotypes from 1016 genotyped F2 progeny, and controlling for genome-wide inbreeding depression, homozygotes for a derived haplotype showed a far lower survival than other genotypes, with only 6.2% of Hn1/Hn1 larvae and 31.3% of the Hn123/Hn123 larvae surviving to the adult stage (GLMM within-family and genotype analyses, Fig. 4A). By contrast, ancestral homozygotes Hn0/Hn0 had a good survival rate (77.6%), and all heterozygous haplotype combinations (Hn0/Hn1; Hn1/Hn123; Hn0/Hn123) displayed similar survival. Inversions therefore harbor fully recessive variants with a strong impact on individual survival in homozygotes. Interestingly, individuals with the Hn1/Hn123 genotype do not experience the deleterious effects of the P_1_inversion (83,8% survival), despite being effectively homozygous for this rearrangement (Fig. 4A). This may indicate that Hn1 and Hn123 harbor different deleterious variants within P_1_, for instance in the region surrounding the gene cortex which shows peaks of differentiation between those two haplotypes (Sup. Fig. S8), or that variants in P_2_ or P_3_ compensate for the deleterious effects of P_1_.

**Fig. 4.**
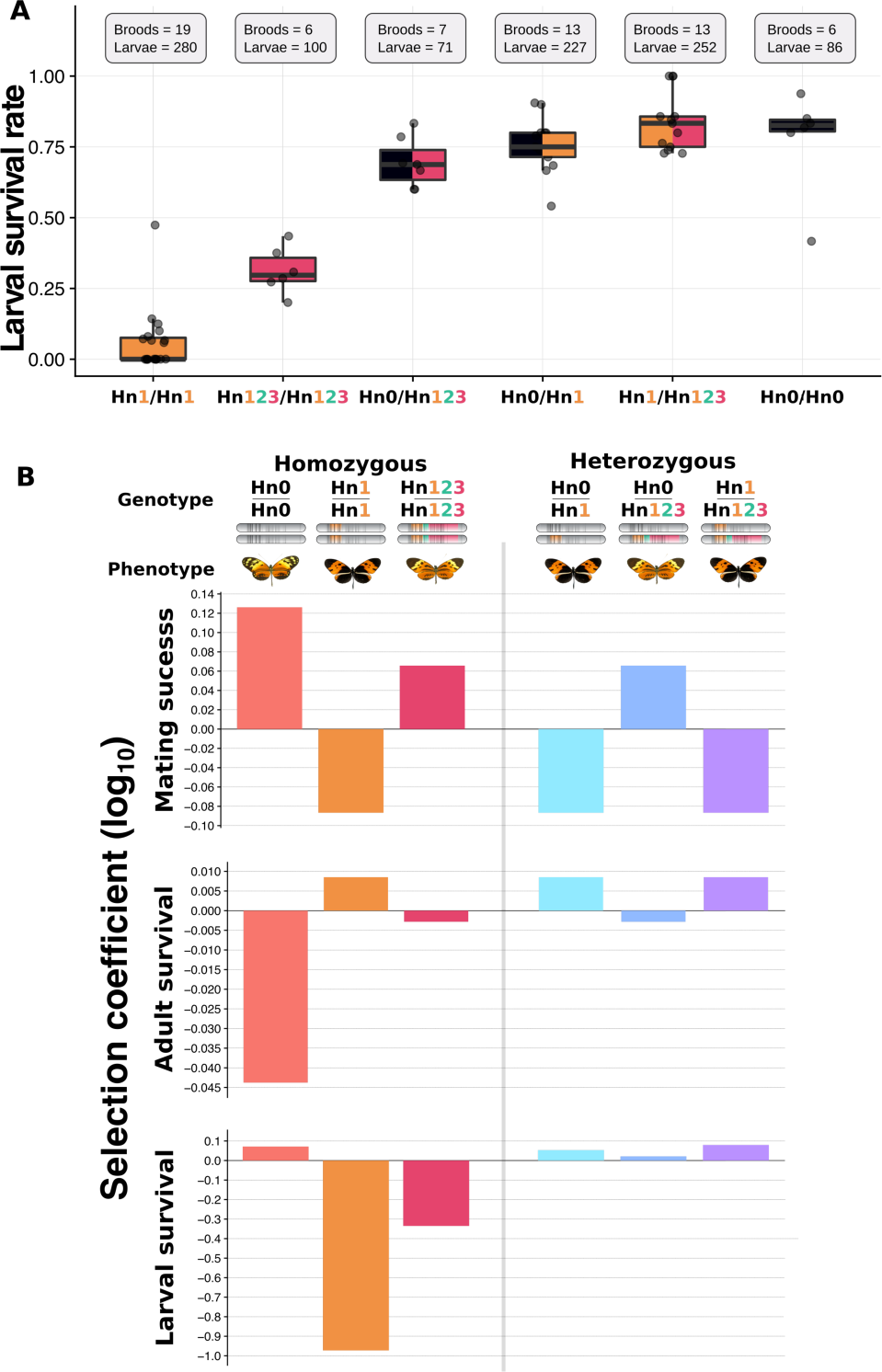
Fitness variation associated with chromosomal inversions at the supergene in *H. numata*. **A.** Larval survival rate for the different supergene genotypes. GLMM analysis confirmed that genotype was a significant predictor of survival (*χ*_2_ = 459.776; df = 5; p<0.001) while experimental cross design was unimportant (*χ*_2_ = 0.8117; df = 2; p = 0.666), validating the joint analysis of all families and crosses. **B.** Variation in fitness components associated with supergene genotypes. Adult survival estimates are based on protection against predator. Selection coefficients were calculated relative to the population mean, and estimated in the *H. numata* population of Tara-poto, Peru. Predation and mating success data come from (16)) and (13)

Inversions have largely been considered for their value in preserving combinations of co-adapted alleles through suppressed recombination with ancestral chromatids, yet this also makes them prone to capturing deleterious mutations (19). Our results bring key insights into how the ecological and genetic components of balancing selection allow inversion polymorphisms to establish. Inversions in *H. numata* show strongly positive dominant effects on adult survival through protection against predators via wing-pattern mimicry, which should lead to their rapid fixation (Fig. 4B, (16)). Yet we found that these inversions are also enriched in deleterious variation from their very formation, as well as from an accumulation of mutations owing to the reduction in recombination-driven purging. The expression of a recessive genetic load associated with inversions inevitably translates into negative frequency-dependent selection (21). The balancing selection acting on these inversions in *H. numata* thus results from their antagonistic ecological and genetic effects: positive selection and dominant effects on adult mimicry but negative frequency-dependent selection through recessive effects on viability (Fig 4B). The initial mutation load associated with the formation and introgression of inversion P_1_ likely initiated the balancing selection as soon as P_1_ rose in frequency, and was further reinforced by the accumulation of deleterious mutations under low recombination. This led to the formation of haplotypes expressing net beneficial effects only when heterozygous.

Individuals carrying inversion P_1_ express disassortative mate preferences, which also balance inversion frequencies in the population (Fig 4B, (13)). Disassortative mating is likely to have evolved in response to the fitness costs associated with homozygous inversions, as selection may have favoured mate preferences minimizing the proportion of homozygous offspring (4). Disassortative mating further hampers the purging of deleterious variation located within the inversions. The initial capture of genetic load in the inversions thus triggered cascading ecological effects and led to the long-term persistence of polymorphism. The low recombination regime associated to inversions also favoured the insertion of transposons, increasing the size of the inverted haplotype. A similar pattern has also been observed in the Papaya neo sex-chromosomes (22) and in the fire ant supergene (23), indicating that this initial increase in size due to accumulation of TE may be a general pattern in the early evolution of polymorphic chromosomes.

Our findings shed new light on the origin and evolution of complex polymorphisms controlled by supergenes and related architectures, such as sex-chromosomes. The build-up of antagonistic fitness effects found here is likely to be a general feature of the formation of inversion polymorphisms and their evolution through time. Therefore, the benefits of structural variants in terms of recombination suppression between ecologically adaptive traits may only explain why they are initially favoured, whereas their maintenance as polymorphisms may be driven by their initial and gradually accumulating mutation load. In summary, balancing selection may not be generated by extrinsic ecological factors, but by intrinsic features of the genetic architecture selected during the evolution of complex phenotypes. Taken together, these novel insights into the consequence of chromosomal rearrangements may explain why inversions are often found polymorphic and linked with complex phenotypes in nature. In a broader context, dissecting the opposing effects of suppressed recombination and how they determine the fate of chromosomal rearrangements may bring new light to our understanding of the variation in genome architecture across the tree of life.

## Methods

### Sampling and sequencing

To investigate the structure of the P supergene allele, we intercrossed wild-caught individuals in cages in order to obtain F2 (or later generation) autozygous individuals (i.e. with the two identical copies of the supergene allele). Samples were either conserved in NaCl saturated DMSO solution at 20°C or snap frozen alive in liquid nitrogen and conserved at −80°C (Sup. Tab. S13). DNA was extracted from the whole butterfly bodies except the head with a protocol adapted from (24), with the following modification. Butterflies were ground in a frozen mortar with liquid nitrogen, 150 mg of tissue powder was mixed with 900µl of preheated buffer and 6µl of RNaseA. Tube were incubated during 120 minutes at 50°C for lysis, and then at −10°C for 10 minutes, with the addition of 300µl of Potassium acetate for the precipitation. One volume of binding buffer was added with 100µl of Serapure beads solution. 3 washing cycles were used and DNA was resuspended in 100µl of EB buffer. Samples 35 and 36 were prepared using the NEBNExt FFPE DNA Repair MIX (NEB). DNA fragment shorter than 20Kb were removed for sample 35 and 36, and shorter than 40kb for samples 26 and 28. 10x Chromium linked-read libraries of 10 autozygous individuals corresponding to 8 different morphs, as well as 2 wild-caught homozygous individuals, were prepared and 2×150bp paired-end reads were sequenced using Illumina HiSeq 2500. Draft genomes were assembled using Supernova (v2.1.1, (25)) (Sup. Tab. S13).

### Whole genome assemblies analysis

The assembled genomes were compared to the *H. melpomene* reference genome (Version 2.5) and to each other using BLAST (26), and LAST (27). Because for some specimens, the supergene was dispersed across multiple scaffolds, we used Ragout2 (28) to re-scaffold their supergene assembly, using as reference the four individual assemblies with the highest quality assembly statistics (n°38, 29, 40, and 26). Genome quality analysis was assessed with BUSCO using the insecta_odb9 database. MAKER (29) was used to annotate the genomes, using protein sequences obtained from the *H. melpomene* genome (v2.5, http://lepbase.org/) in combination with an *H. numata* transcriptome dataset (30). RepeatModeler (31) was used to identify unannotated TEs in the 12 *H. numata* genomes. Unknown repeat elements detected by RepeatModeler were compared by BLAST (26) (-evalue cut-off 1e^-10^) to a transposase database (Tpases080212) from (32). TE identified were merged with the Heliconius repeat database (Lavoie et al. 2013) and redundancy was filtered using CDHIT (REF) with a 80 % identity threshold. Repeat-Masker (31) was then used to annotate transposable elements and repeats using this combined database and results were parsed with scripts from https://github.com/4ureliek/Parsing-RepeatMasker-Outputs.git.

### Population Genomic Analysis

Whole genome re-sequence data from *H. numata* and other Heliconius species from (17) were used, as well as 37 new wild-caught *H. numata* specimens. For the latter samples, butterfly bodies were conserved in NaCl saturated DMSO solution at −20°C and DNA was extracted using QIAGEN DNeasy blood and tissue kits according to the manufacturer’s instructions with RNase treatment. Illumina Truseq paired-end whole genome libraries were prepared and 2×100bp reads were sequenced on the Illumina HiSeq 2000 platform. Reads were mapped to the *H. melpomene* Hmel2 reference genome (33) using Stampy (version 1.0.28; (34)) with default settings except for the substitution rate which was set to 0.05 to allow for expected divergence from the reference. Alignment file manipulations were performed using SAMtools v0.1.3 (35). After mapping, duplicate reads were excluded using the MarkDuplicates tool in Picard (v1.1125; http://broadinstitute.github.io/picard) and local indel realignment using IndelRealigner was performed with GATK(v3.5; (36)). Invariant and polymorphic sites were called with GATK HaplotypeCaller, with options – min_base_quality_score 25 –min_mapping_quality_score 25 -stand_emit_conf 20 –heterozygosity 0.015. VCF data were processed using bcftools (37). PCA analyses were computed with the SNPRelate R package (38), using 5kb windows. Using Phylobayes (39), on 5kb sliding windows, we estimated the most recent coalescence event between Hn0+Hn1 and Hn123, which corresponds to age of the last recombination between Hn0+Hn1 and Hn123, and 2) the time to the most recent common ancestor (TMRCA) of all Hn123 haplotypes. This provides respectively the upper (1) and the lower (2) bounds of the date of the inversion event (Sup Fig. S7). In order to compute the Fst and standard population genetic analyses, we manually curated the phasing of heterozygous individuals since computational phasing packages such as SHAPEIT or BEAGLE were found to introduce frequent phase switch errors. For each heterozygous SNP in inversion regions, if one and only one of the two alleles is observed in more than 80 % of individuals without inversions (Hn0), this allele is considered as being on the haplotype 1, the other being on haplotype 2. For SNPs which did not fit this criterion, each allele was placed randomly on one of the two haplotypes.

### Deleterious mutation accumulation

SnpEff (40) with default was used to annotate the *H. numata* SNPs using the *H. melpomene* reference genome annotation. We computed the ratio of synonymous and non-synonymous variants (pN/pS), the rate of synonymous and non-synonymous substitution (dN/dS) compared to H. melpomene, and the direction of selection with DoS = Dn/(Dn + Ds) Pn/(Pn + Ps) (20), using all individuals, or only those homozygous for a given inversion type, for every genes larger 5kb (to ensure there is a several SNP within each gene). Whole genome distribution was computed on 500kb non-overlapping sliding windows.

### Fitness Assay

*H. numata* specimens used for the fitness analyses originated the Tarapoto valley, San Martin, Peru. Brood designs are illustrated in Sup. Fig. 10. First, F1 P heterozygotes butterflies were generated by crossing F0 wild males to captive bred virgin females. Unrelated F1 male-female pairs were then selected for their P genotype and hand paired to generate an F2 progeny. We specifically designed these crosses to generate a F2 progeny containing both homozygotes and heterozygotes, within a single family. Larvae were monitored twice a day to assess survival or mortality. Upon death or butterfly emergence, individuals were stored in 96° ethanol until genotyping. We generated a total of 486 F2 progeny from 6 independent replicate of broods for the F1 Hn0/Hn1 x Hn0/Hn1 cross, 504 F2 progeny from 6 brood of the F1 Hn1/Hn123 x Hn1/Hn123 cross and 454 F2 progeny from 7 broods of the F1 Hn1/Hn123 x Hn0/Hn1 cross. Supergene genotypes was assessed using (13) methodology. Briefly, the amplification of the Heliconius numata orthologue of HM00025 (cortex) (Genbank accension FP236845.2), included in the supergene P enables to discriminate between the distinct supergene haplotype by PCR product size: Hn1 (∼1200bp), Hn123 (∼800bp) and Hn0 (∼600 bp). 1,016 F2 progeny could be genotyped. For each of the 19 broods, we used a Chi-squared test of independence to assess variation in survival between the different genotypes of the F2 progeny. When significant, the Freeman-Tukey deviates (FT) was compared to an alpha = 0.05 criterion, and corrected for multiple comparisons using the Bonferroni correction. To compare genotype survival between families and crosses we performed generalized linear mixed models analysis followed by a Tukey’s HSD post-hoc test (package “lme4” (41); in R version 3.1.3, (42)), with the survival of an individual with a given genotype as the response variable (binomial response with logit link). The significance of the predictors was tested using likelihood ratio tests. The genotype was a covariate predictor, crosses was a fixed effect and family identity as a random effect to control for non-independence of measures. Plots were created with ggplot2 (43).

## ACKNOWLEDGEMENTS

We thank Emmanuelle d’Alençon and Marie-Pierre Dubois for their help in the lab, Thomas Aubier for being a DNA extraction wizard, Melanie McClure, Mario Tuatama, Ronald Mori-Pezo for their help during field work, Patrice David for his careful and critical reading of the manuscript, Konrad Lhose, Dominik Laetsch, Benoit Nabolz, Pierre-Alexandre Gagnaire, Mathieu Gauthier, Claire Lemaitre, Fabrice Legeai and Anna-Sophie Fiston-Lavier for insightful discussions. We thank the Peruvian government for providing the necessary research permits (236-2012-AG-DGFFS-DGEFFS, 201-2013-MINAGRI-DGFFS/DGEFFS and 002-2015-SERFOR-DGGSPFFS). This research was supported by Agence Nationale de la Recherche (ANR) grants ANR-12-JSV7-0005 and ANR-18-CE02-0019-01 and European Research Council grant ERC-StG-243179 to MJ and by fellowships from the Natural Sciences and Engineering Research Council of Canada and a Marie Sklodowska-Curie fellowship (FITINV, N 655857) to MC. This project benefited from the Montpellier Bioinformatics Biodiversity platform supported by the LabEx CeMEB, ANR “Investissements d’avenir” program ANR-10-LABX-04-01. MGX acknowledges financial support from France Génomique National infrastructure, funded as part of ANR “Investissement d’avenir” program ANR-10-INBS-09.

## AUTHOR CONTRIBUTIONS

P.J., M.C., and M.J. designed the study. P.J., M.C., A.W., and M.J. wrote the paper. P.J., A.W., and M.J. generated the genomic data. M.C., H.B and V.L. generated the RNAseq data. P.J. performed the genomic analyses with input from A.W.. M.C. managed butterfly rearing and performed fitness assays. H.P. performed whole genomes sequencing. All authors contributed to editing the manuscript.

## COMPETING FINANCIAL INTERESTS

The authors declare no competing interests

## Supplementary Note 1: Figures

**Fig. S1.**
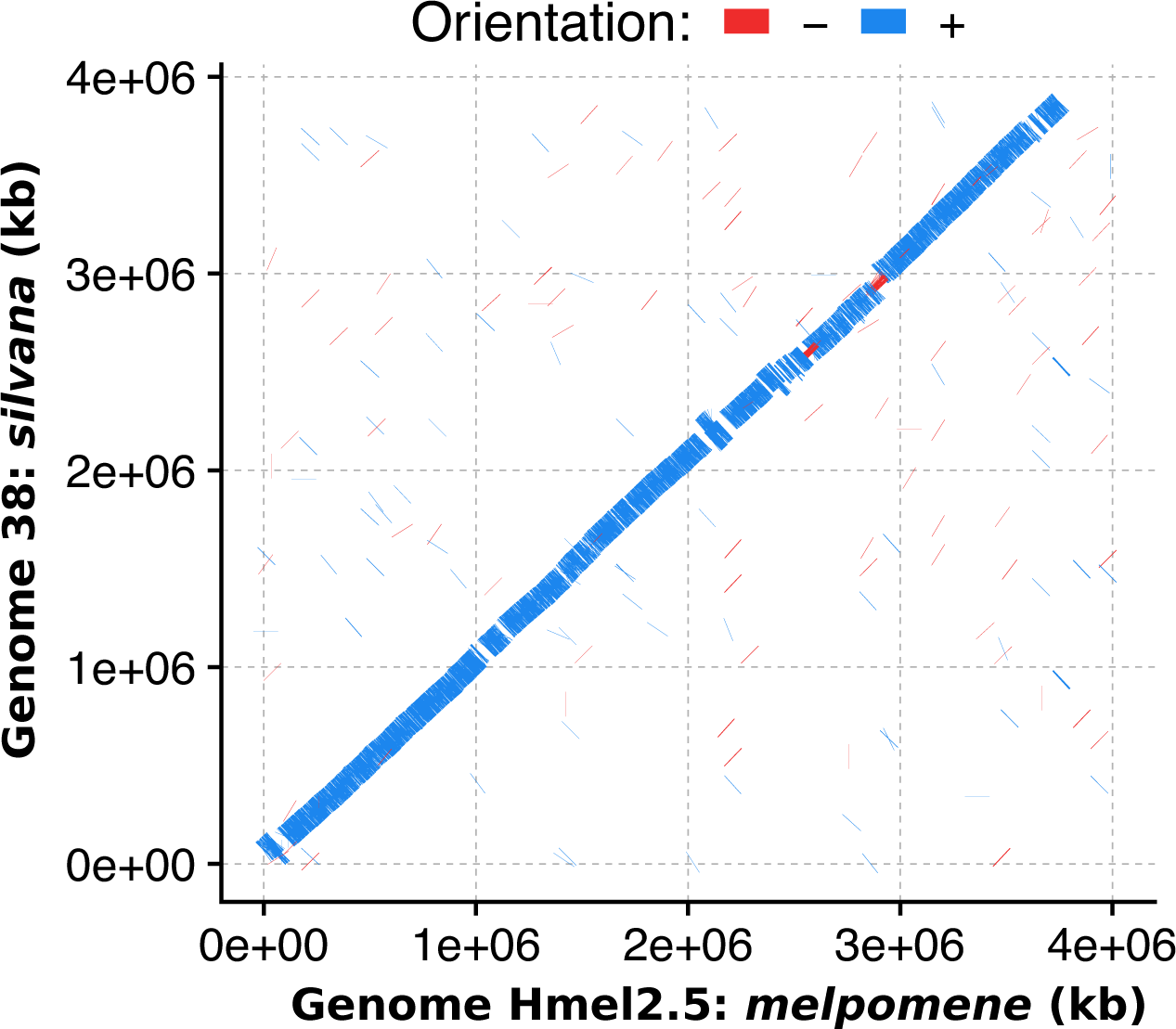
Alignment of genome assemblies of *H. numata silvana* (Hn0, genome 38) and the *H. melpomene* reference genome (Hmel2.5) focused on the region of the supergene on chromosome 15. No major chromosomal rearrangements are observed between Hn.0 and *Heliconius melpomene* on chromosome 15.

**Fig. S2.**
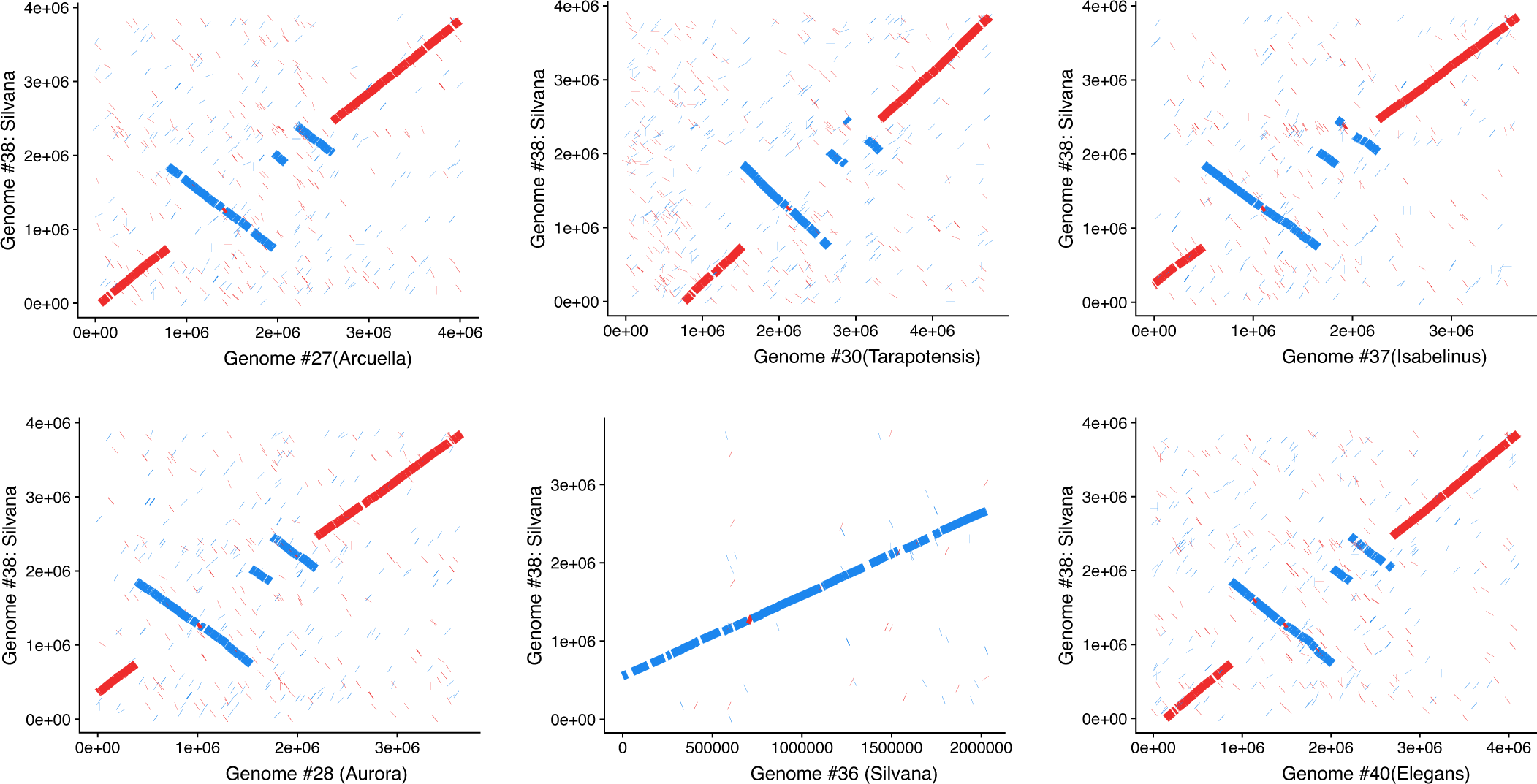
Alignment of the supergene region of genome 38 (.*H. n. silvana*) against other *H. numata* genome assemblies.

**Fig. S3.**
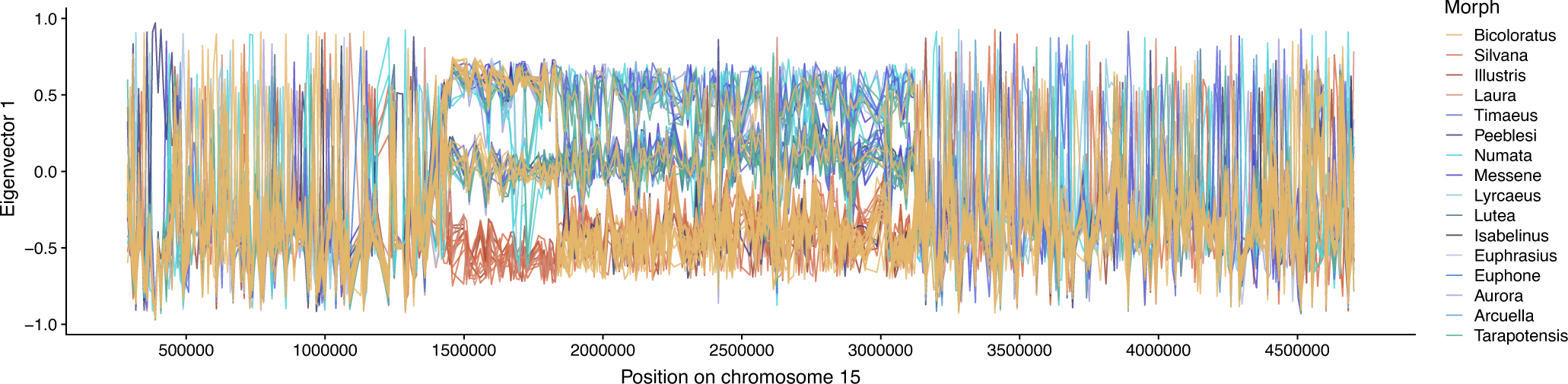
Sliding window PCA computed along the supergene for all specimens. Computed on 5kb sliding windows. Each line represents the position of a specimen on the first axis of the PCA along chromosome 15. See Sup. Fig. 4S for summary PCAs, not computed on sliding wind. ows but on the whole regions.

**Fig. S4.**
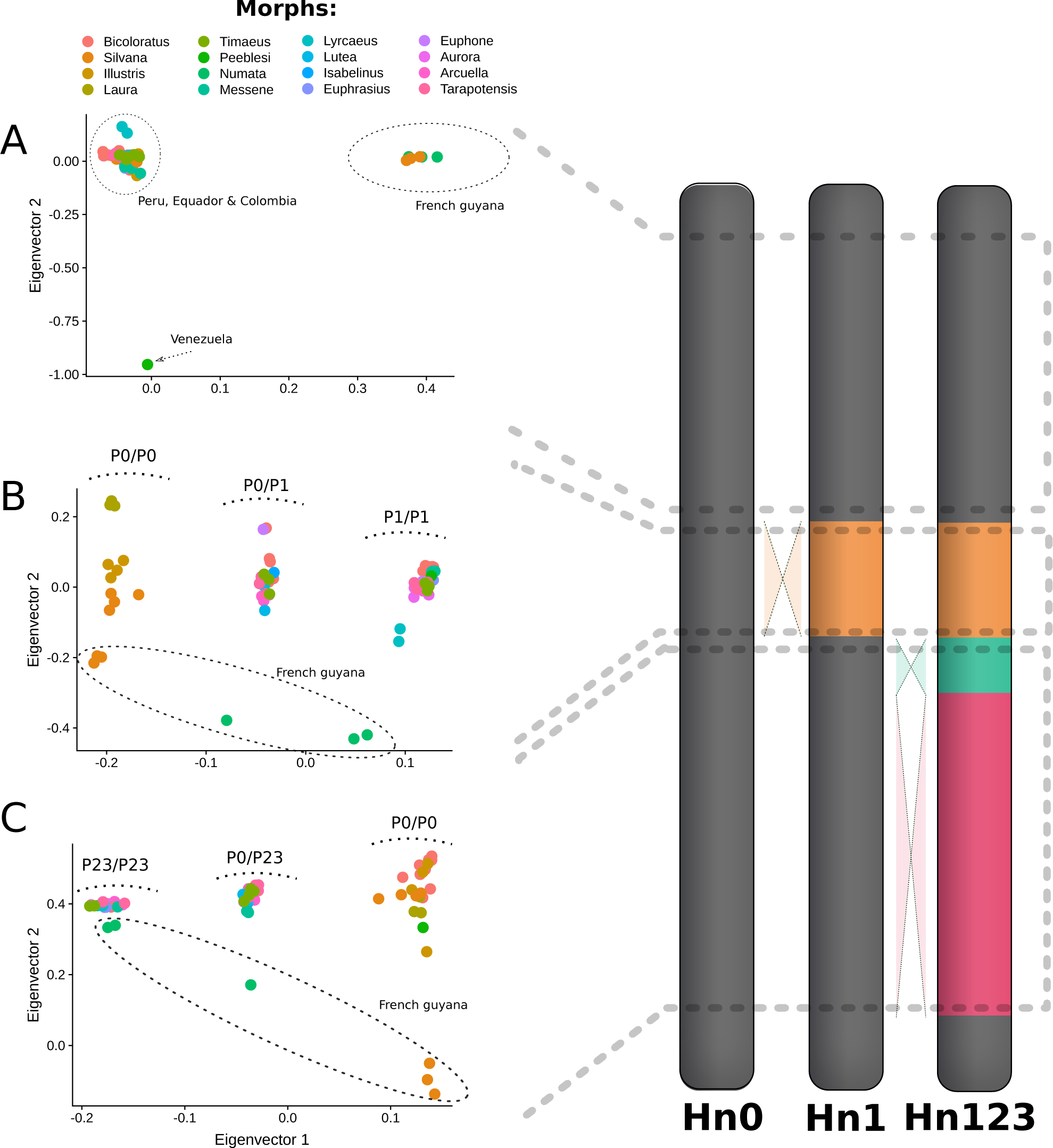
PCA computed on SNPs, on the inversion segments and outside the supergene, for all specimens. Each dot represents the position of a specimen on the PCA two first axis. **A.** PCA computed on SNPs on the chromosome 15 but not within the supergene region. The PCA reflect the geographic structure of the dataset. **B.** PCA computed on SNPs on P_1_ segment. The first axis of the PCA reflects individual genotypes for the inversion : homozygote for the ancestral gene order (P0/P0), Homozygote for the inversion (P1/P1), or heterozygote (P0/P1). The second axis of the PCA reflects the geographic structure of the dataset. **C.** PCA computed on SNPs on P_2_+P3 segment. The first axis of the PCA reflects individual genotypes for the two inversions : homozygote for the ancestral gene order (P0/P0), Homozygote for the two inversions (P23/P23), or heterozygote (P0/P23); the second axis of the PCA reflects the geographic structure of the dataset.

**Fig. S5.**
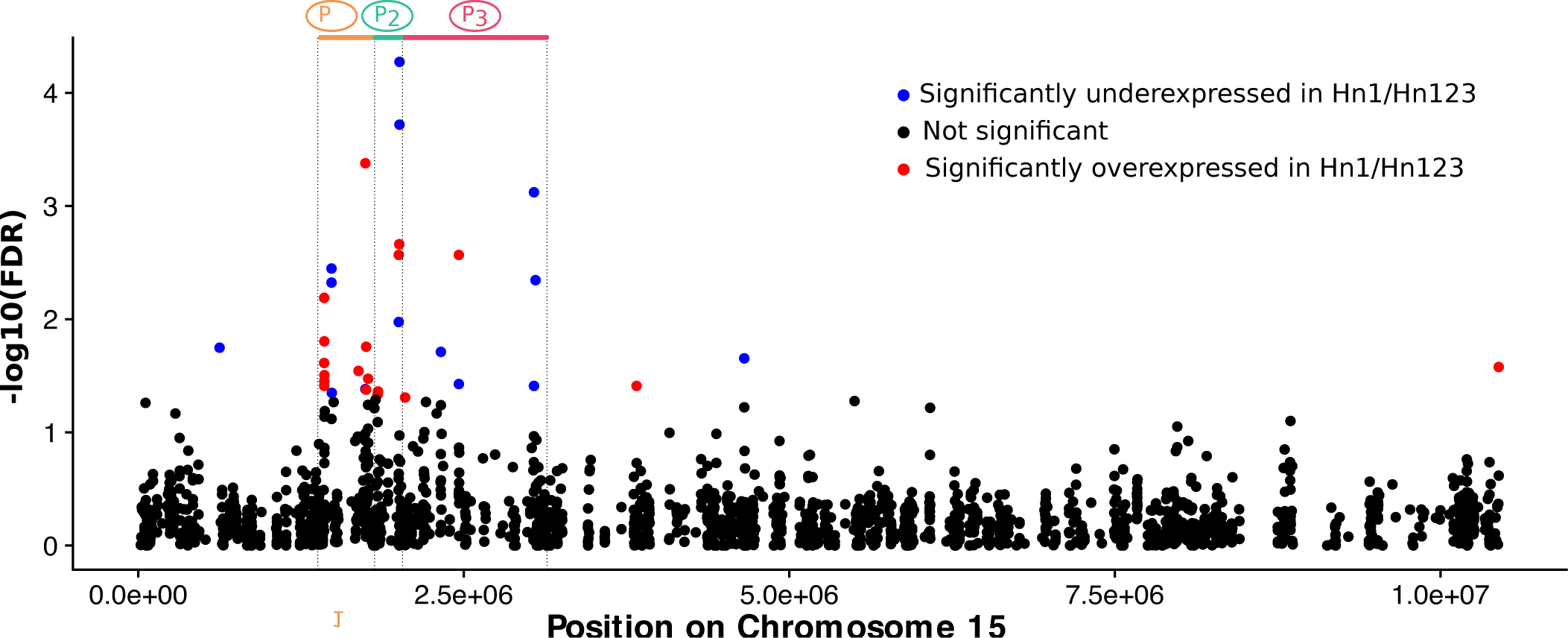
Differential gene expression across the chromosome 15. Expression difference in early pupal (24h) wing discs between Hn0 and Hn1/Hn123. RNAseq data from (1) were reanalysed using the EdgeR R package (2)). The -log10 of the false discovery rate is plotted along the chromosome 15, with each dot representing a different transcript, and reveal that genes within the inversion segments are differentially expressed between Hn0 and Hn1-Hn123.

**Fig. S6.**
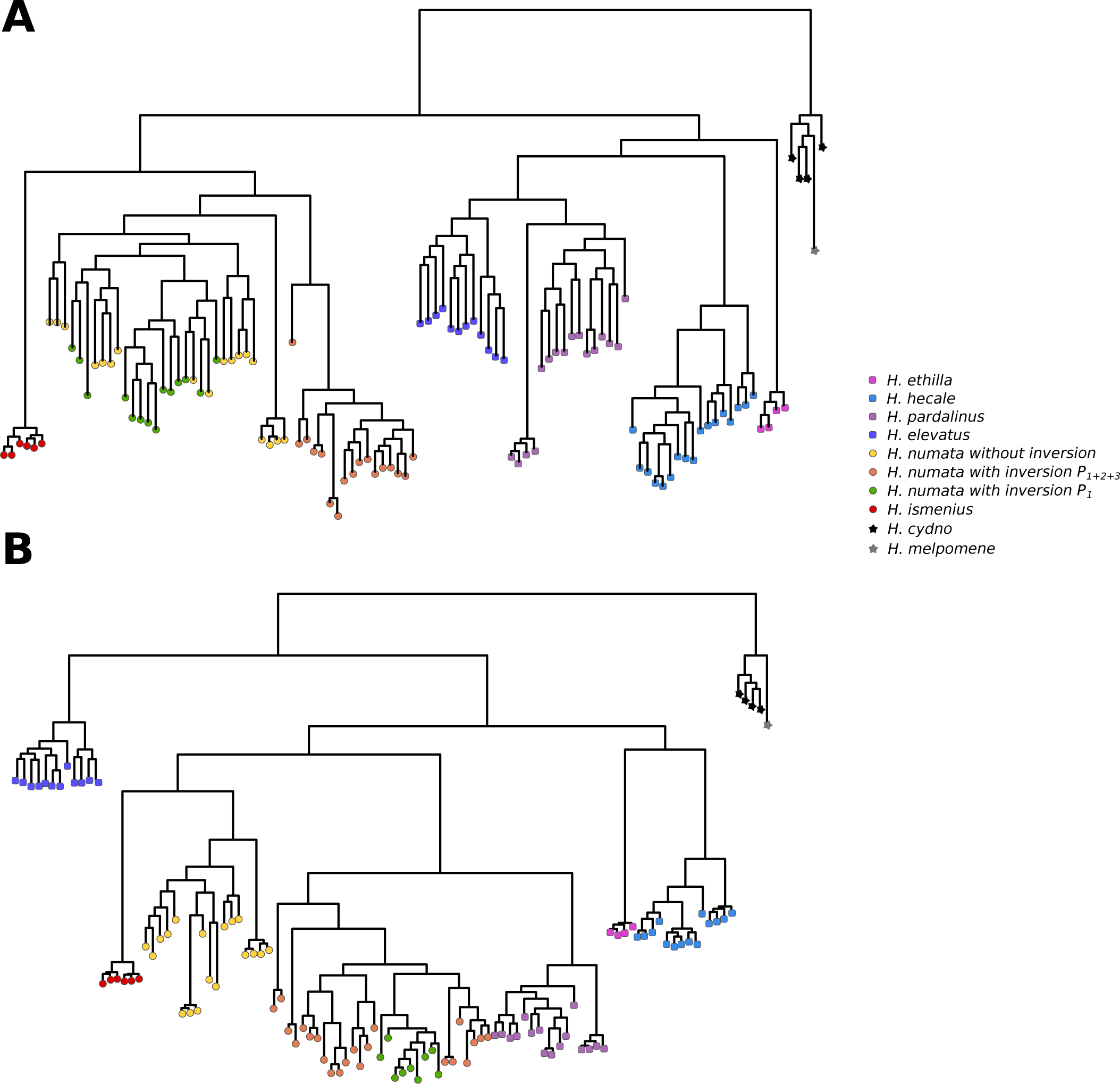
Phylogenies of the silvaniform clade with *H. cydno* and *H. melpomene* as outgroups, using the genomic segments orthologous to P_1_, P_2_ and P_3_ in *H. numata*. Phylogenies computed with RAxML (3) using the GTRCAT model and only individuals homozygous for the inversions or the standard arrangement. **A.** Phylogeny of segments orthologous to P_2_ and P_3_. This shows the unique origin of the P_2_ and P_3_ inversions within *H. numata*. **B.** Phylogeny of segments orthologous to P_1_. This show the introgression of P_1_ from *H. pardalinus* into *H. numata*. Incongruent position of *H. elevatus, H. hecale* and *H. ethilla* result from incomplete lineage sorting at the clade level around the gene cortex and to gene flow among species of the clade (especially an introgression between *H. elevatus* and *H. melpomene*) (4).

**Fig. S7.**
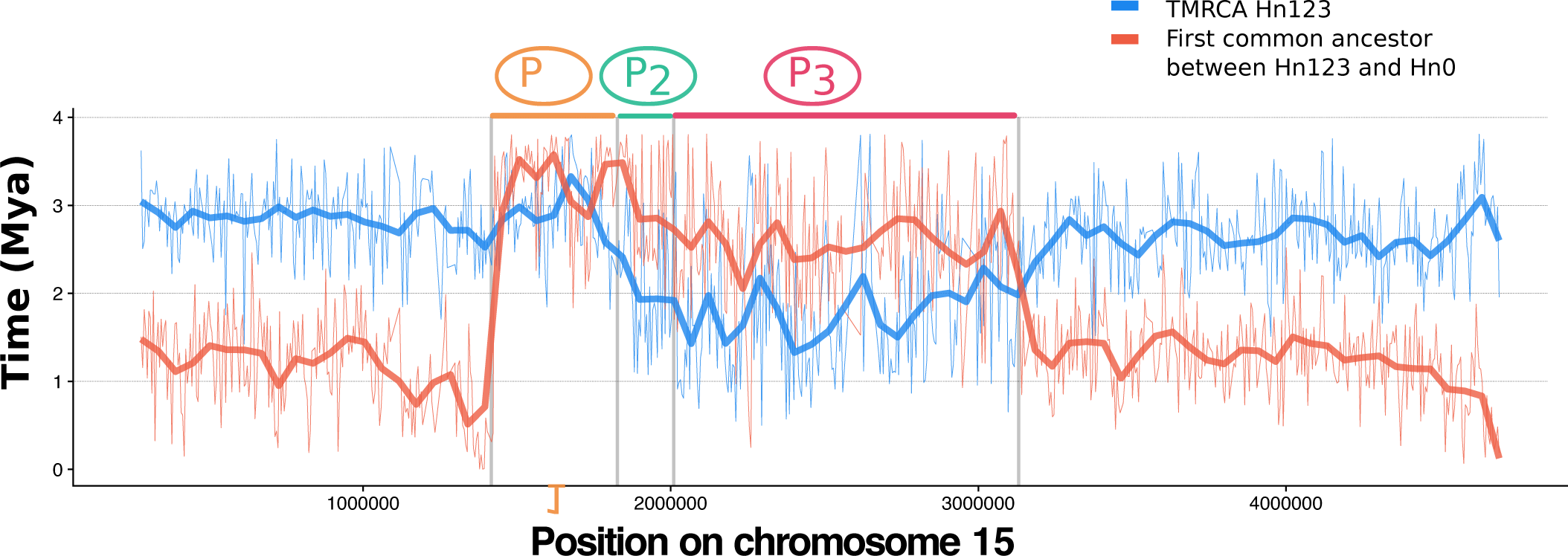
Analysis of divergence times between Hn123 and Hn0 along the chromosome 15. Divergence time estimates computed with Phylobayes on 5kb sliding windows. Bold red and blue lines represent the LOESS smoothing (span = 0.05) of the raw data (thin lines) and give the upper and lower bound of the times inversions P_2_ and P_3_ occurred. This supports the formation of P supergene by the stepwise accretion of P_1_, P_2_ a.nd P_3_

**Fig. S8.**
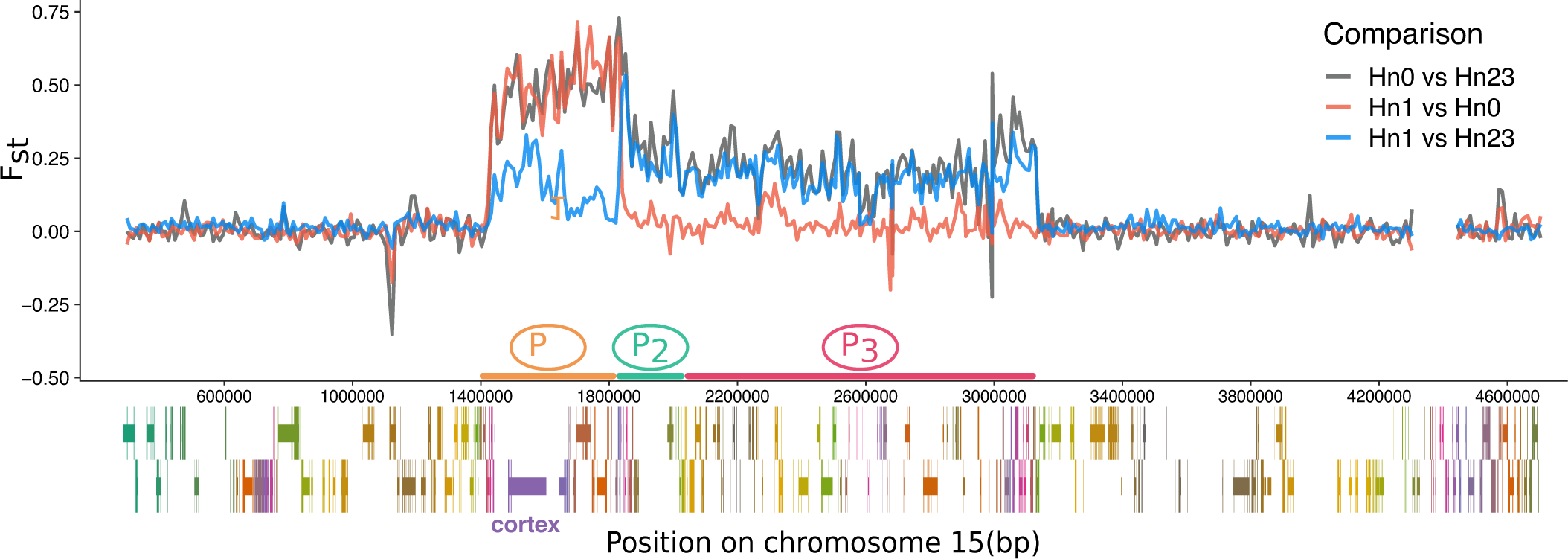
Fst analysis between the three main supergene alleles : without inversion (Hn0), with P_1_ inversion (Hn1) and with all three inversion P_1_,P2 and P_3_ (Hn123) A “suspension bridge” pattern of differentiation can be observed at P_2_-P3 by comparing Hn123 to Hn0 and Hn1 haplotypes, suggesting the rare occurrence of recombination around the center of the inversion, as predicted by (5). A peak of differentiation can be seen between Hn1 and Hn123 around the gene cortex, which controls melanic variations of the wing pattern in Heliconius butterflies (6). This peak was unexpected since these two classes of haplotypes have the same genomic orientation (P1 inversion) in this region. Moreover, this region also show the highest differential gene expression when comparing Hn1 to Hn123 (Sup. Fig. S5). Analyses of assemblies as well as of read coverage (data not shown) do not support the presence of major rearrangements between Hn1 and Hn123 at this position, suggesting that this peak of differentiation on cortex is caused by selection on wing pattern divergence rather than recombination suppression via structural variation.

**Fig. S9.**
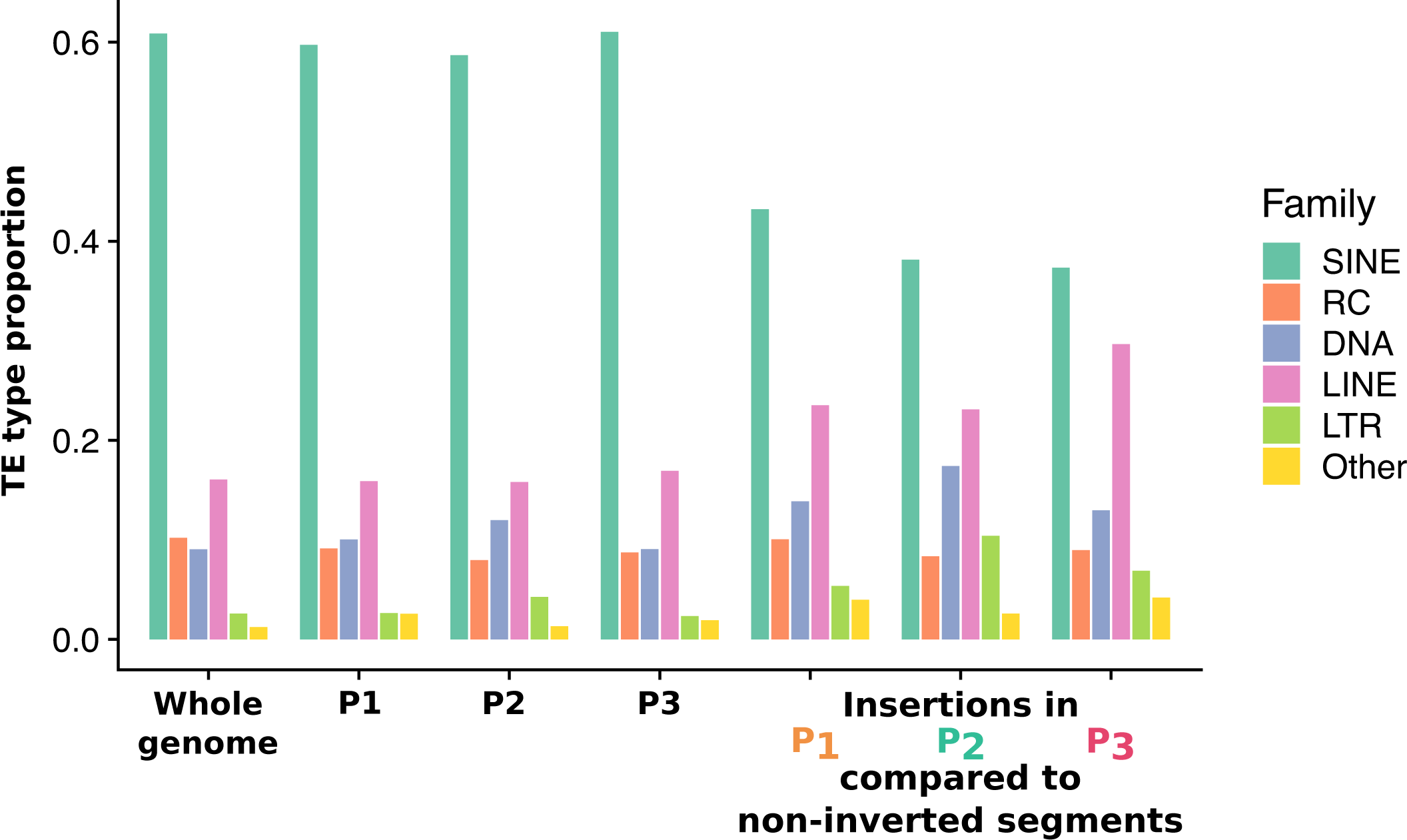
Proportion of TE classes in whole genome, inversions, and insertions in inversions.

**Fig. S10.**
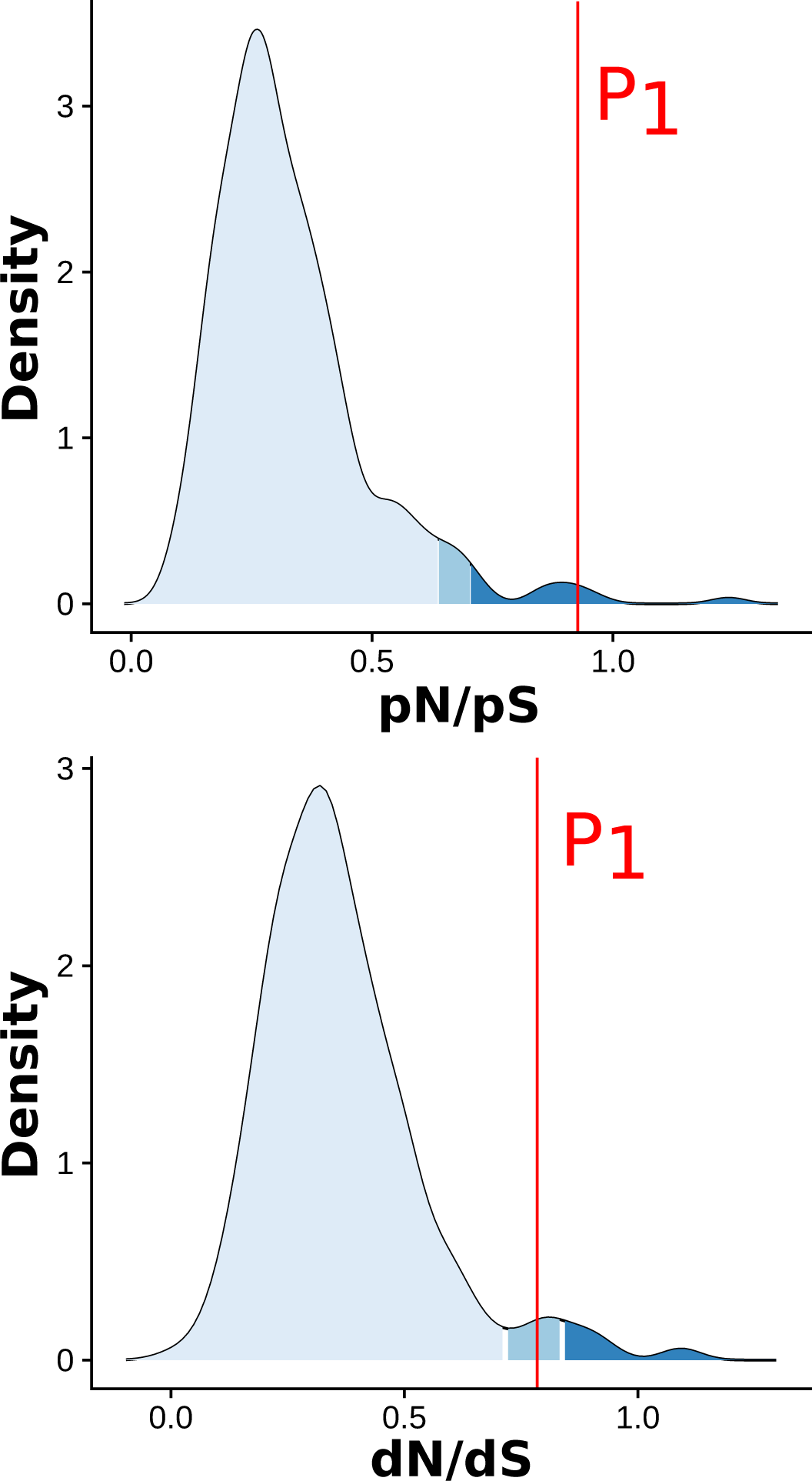
Mutation accumulation analysis on *H. pardalinus*. Density curve representing the whole genome distribution computed on 500kb windows across 12 *H. pardalinus* specimens. P_1_ shows an increase in non-synonymous polymorphisms and substitutions c. ompared to whole genome.

**Fig. S11.**
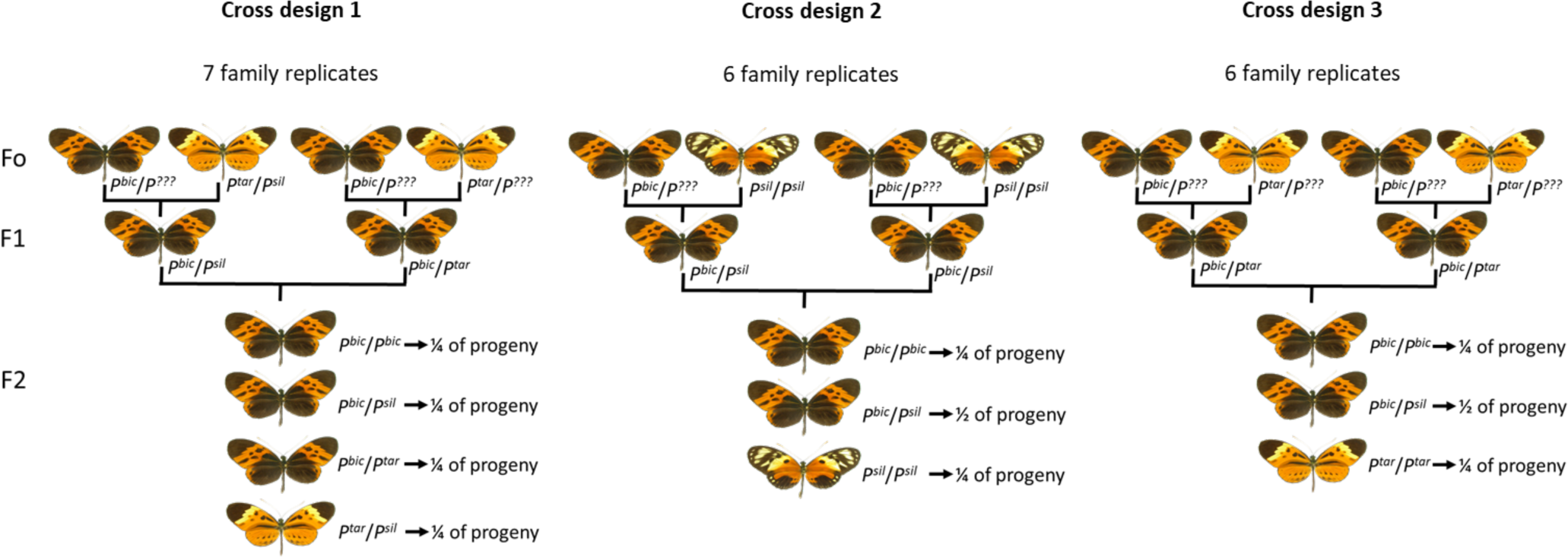
Experimental crosses designed to assess the survival of the larvae of the distinct genotypes at the supergene P.

## Supplementary Note 2: Tables

**Fig. S12.**
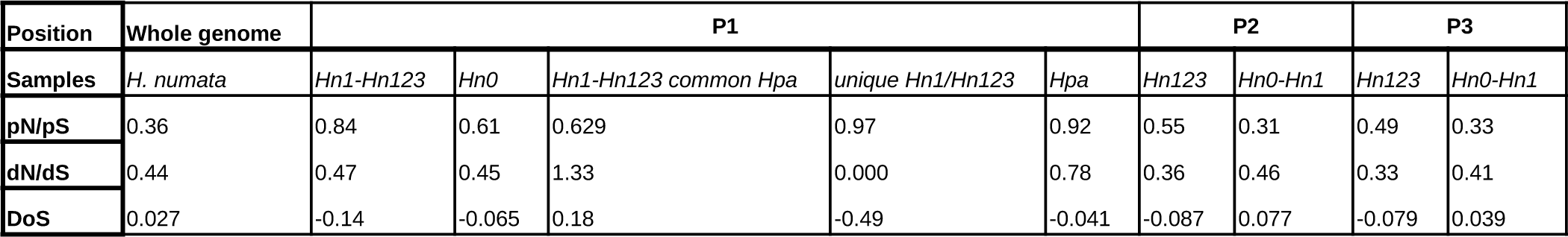
Accumulation of deleterious variants in inversions. dN/dS, pN/pS and Direction of Selection computed on the whole genome or only on segments P_1_, P_2_, or P_3_. Only samples homozygotes for the ancestral or the inverted gene order are used for the analysis. Hn0 display the ancestral gene order at P_1_, P, and P_3_. Hn1 are inverted at P_1_ and non-inverted at P_2_ and P_3_. Hn123 are inverted at P_1_, P_2_, and P_3_. Because P_1_ was introgressed from *H. pardalinus* (Hpa), we were able estimate parameters on mutations that are unique to Hn1-Hn123, which occurred after the inversion formation, and on mutations that are common to Hn1-Hn123 and Hpa, which occurred before the introgression. Inverted segments consistently show a more negative direction of selection compared to non-inverted segments and a higher pN/pS ratio, suggesting a lower efficiency of selection to purge deleterious variants in inversion. Contrarly, dN/ds ratio are slightly lower in inverted compared to non-inverted segments. P_1_ segments help to understand this pattern. Non-synonymous SNPs that occurred in coding region of P_1_ in Hpa before the introgression (“Hn1-Hn123 common Hpa”) underwent a very high rate of fixation in Hn1-Hn123 (dN/dS=1.33), but none of the SNPs that occurred in Hn1-Hn123 after the introgression is fixed (dN/dS=0,000). This suggest that the indermediate dN/dS values observed at inversions may result from the balance between the very high rate of fixation during inversions formation (and introgression) and the reduction of fixation rate during their su.bsequent evolution, likelly because of recombination suppression.

**Fig. S13.**
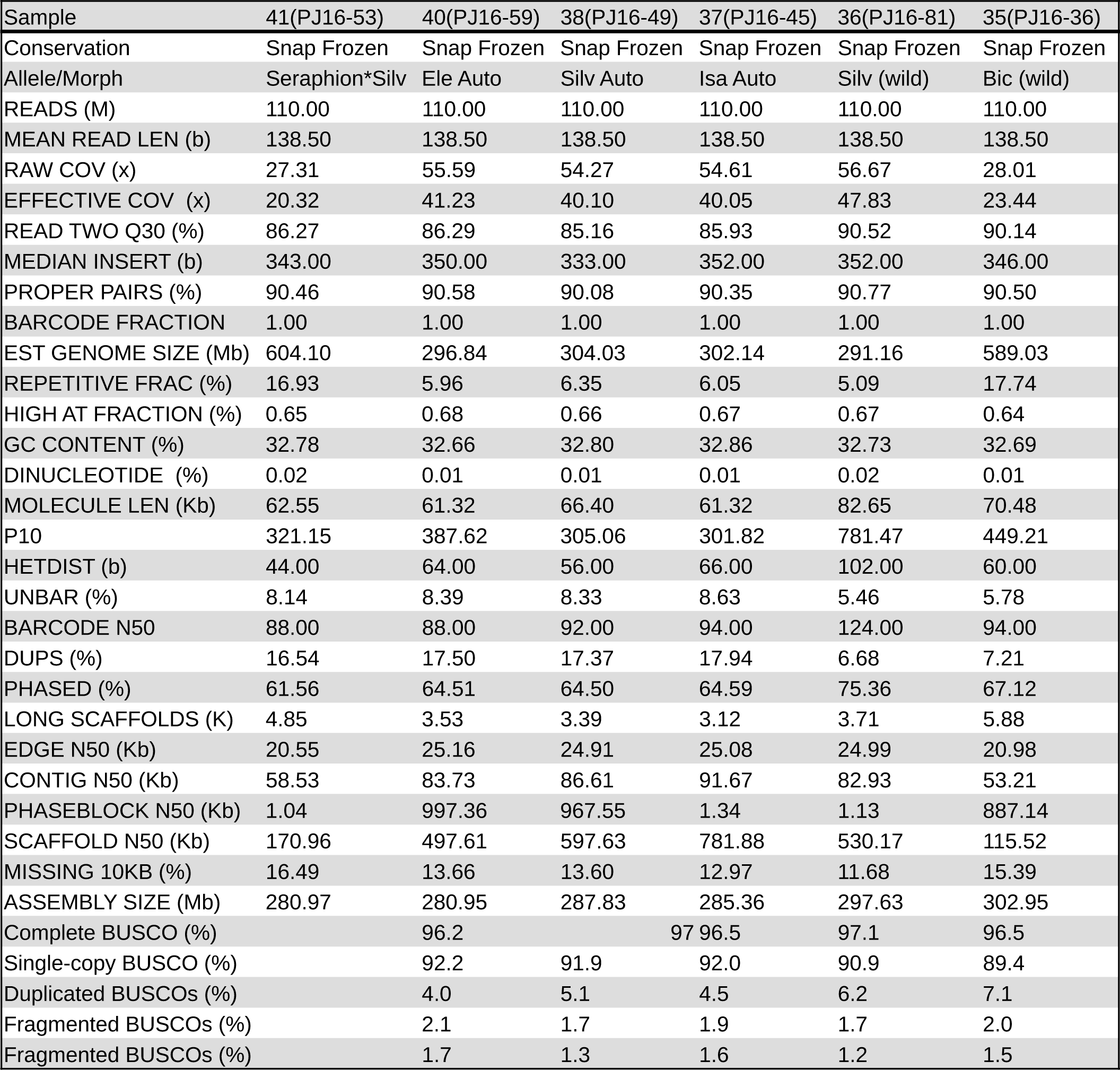
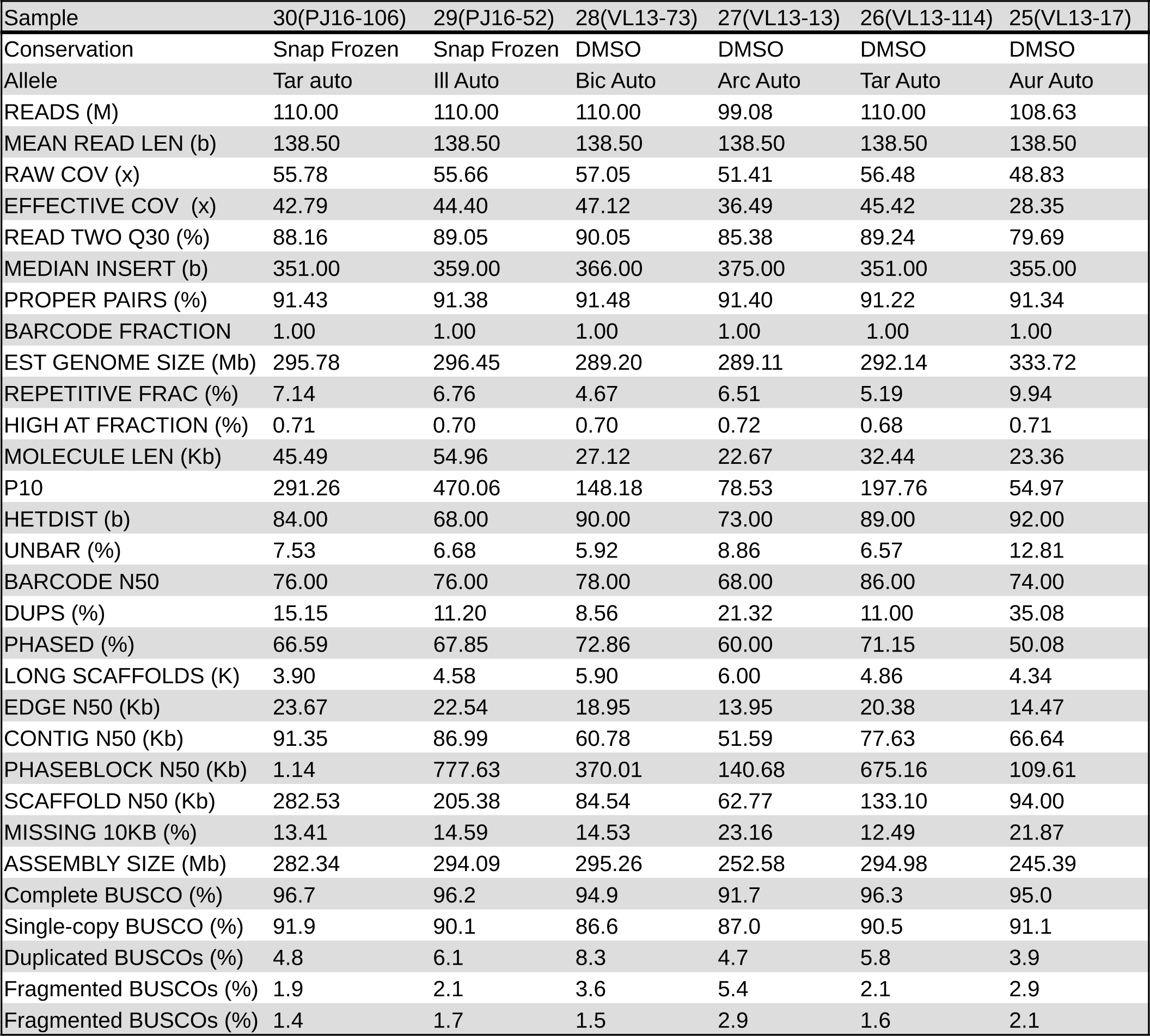
Summary of genome assemblies quality.

